# Lipid nanoparticle delivered intrabodies for inhibiting necroptosis and pyroptosis

**DOI:** 10.1101/2025.05.06.652432

**Authors:** Veerasikku Gopal Deepagan, Xiuquan Ma, Farzaneh Bazregari, Jiyi Pang, Jan Schaefer, Joanne M. Hildebrand, Ruby K. Dempsey, Marcel Doerflinger, Christopher A. Baldwin, Florian I. Schmidt, James M. Murphy, Ranja Salvamoser, James E. Vince

## Abstract

Intrabodies are intracellularly expressed high-affinity protein binders such as nanobodies and monobodies that offer an alternative approach to small molecules. However, the maturation of intrabody technology into new therapeutic modalities has been limited by the availability of a clinically relevant delivery system enabling sufficiently high levels of protein to be expressed in the cytosol. Here, we use lipid nanoparticle (LNP) systems based on clinically approved formulations for the efficient intracellular delivery of mRNAs encoding for intrabodies targeting mixed lineage kinase domain-like pseudokinase (MLKL) and apoptosis-associated speck-like protein containing a CARD (ASC), key mediators of the necrotic cell death modalities, necroptosis and pyroptosis, respectively. LNP delivery of intrabody mRNA resulted in robust protein expression, with a MLKL binding intrabody preventing MLKL membrane translocation and protecting against necroptotic cell death. Similarly, LNP delivery of a bivalent intrabody targeting the inflammasome adaptor protein ASC protected against NLRP3 and AIM2 inflammasome-driven responses, including caspase-1 and IL-1β activation and gasdermin D-driven pyroptotic killing. These findings establish that LNPs harbouring anti-necrotic intrabody mRNAs allow for sufficient intracellular expression to neutralize necrotic cell death signalling and provide a general, clinically relevant, strategy for delivering therapeutic intrabodies into cells.

## Introduction

Antibody therapeutics are widely used to treat a broad spectrum of diseases, including cancer, autoimmune disorders, and infectious diseases [1, 2]. However, their application can be limited by several factors, including side effects such as acute anaphylactic reactions and cytokine release syndrome, poor tissue penetration due to their large size (∼150 kDa), and high production costs [3]. Consequently, alternative antigen binding scaffolds such as nanobodies (single-domain antibody fragments derived from camelids) and monobodies (based on the type III fibronectin domain) have been explored, with nanobody-based therapies targeting extracellular proteins, such TNF, PD-L1 and von Willebrand factor approved for clinical use, and many others undergoing clinical evaluation [4, 5].

When compared to conventional antibodies, attractive unique features of nanobodies, monobodies and other protein binders (*e*.*g*., designed ankyrin repeat proteins) are their small size of ∼ 15 kDa enabling access to “hidden” epitopes, simple single-domain structure (*i*.*e*., can be encoded by a single DNA or mRNA), lack of a Fc region (thereby avoiding the potential for triggering anaphylaxis), and their stability in reducing environments [4, 5]. These features also mean that single domain antibodies and antibody mimetics can fold properly inside cells to bind, with high affinities, their targets, resulting in the term “intrabodies” to collectively describe their ability to function intracellularly [6]. Intrabodies have been extensively used as research tools, such as in live-cell imaging and as biosensors and to help uncover new biology, although their capacity to inhibit proteins classified as undruggable highlights their potential as a new class of therapeutics [7]. For example, studies have demonstrated how intrabodies can block the signalling capacities of hard to drug targets, such as the cancer drivers RAS [8], β-catenin [9] and LMO2 [10], or key molecules required for programmed necrotic cell death pathways, necroptosis [11] and pyroptosis [12, 13], implicated in numerous inflammatory conditions [14]. However, although a number of intrabody transporting systems, such as micro-injection, lentiviral transduction, and the incorporation of cell-penetrating motifs, have been investigated [15], clinically proven modalities for the delivery of intrabodies into cells remains a major challenge for taking them forward as viable drugs.

Lipid nanoparticles (LNPs) represent a promising strategy for intracellular delivery of nucleic acids, including mRNA. These nanometre-sized, self-assembled vesicles protect mRNA from temperature variation, extracellular RNase degradation, and premature clearance by the mononuclear phagocyte system or renal filtration [16, 17]. LNPs safely transport mRNA across physiological barriers, predominantly via endocytosis, and deliver the payload into the cytosol [17]. Moreover, LNPs can be engineered for tissue-specific delivery by altering physicochemical properties such as lipid composition [18], surface charge [19], shape [16], particle size [20] and tagging with targeting ligands [21]. These modifications can enable LNP tissue and cellular trophism, thereby minimizing off-target effects. In addition, improved *in vitro* transcription methods with optimized untranslated regions (UTRs), modified bases, and effective 5′-cap structures have contributed to making LNP-mRNA technology a powerful tool for intracellular protein delivery that is bringing several new biological medicines into the clinic [16].

Necroptosis and pyroptosis are two genetically encoded necrotic cell death pathways that evolved to counter pathogen infections, but their dysregulation coupled with their inflammatory nature have implicated them in driving numerous infectious, inflammatory and autoimmune diseases [14, 22-24]. MLKL is the terminal effector of necroptosis, and when activated following death receptor, toll-like receptor (TLR) or Z-DNA binding protein 1 (ZBP1) signalling, it oligomerizes and unleashes the 4-helical bundle (4HB) domain, which then interacts with negatively charged membrane lipids to cause plasma membrane lysis and the release of immunogenic intracellular contents [14, 25-27]. On the other hand, pyroptosis is initiated canonically through cytosolic pattern recognition receptors (PRRs) such as NLRP3, a sensor of cellular stress and potassium ion efflux, AIM2, a sensor of DNA, and Pyrin, an indirect sensor of bacterial toxins. When triggered, these inflammasome sensor proteins recruit the adaptor ASC which binds caspase-1. Subsequently, proximity-induced activation of caspase-1, for which ASC is essential [28], results in caspase-1 maturation of the inflammatory cytokines IL-1β and IL-18, and also processing of gasdermin D (GSDMD) [29, 30]. The N-terminal domain of GSDMD subsequently forms plasma membrane pores, resulting in pyroptotic cell death and enabling the efficient release of activated IL-1β and IL-18 [31].

Despite the promise of targeting necroptosis or pyroptosis to treat diverse inflammatory-associated diseases, to date, no intracellular anti-necrotic drug has progressed beyond clinical trials. Here, we sought to leverage clinically approved LNP-mRNA technology to deliver intrabodies targeting two key mediators of necrotic cell death – an intrabody that targets MLKL, terminal effector of necroptosis, and an intrabody targeting ASC, the common adaptor protein essential for pyroptosis downstream of inflammasome sensor protein activation. We demonstrate that LNP delivery of MLKL and ASC intrabody mRNA results in stable protein expression and potently blocks necroptosis and pyroptosis, respectively, paving the way for future trials examining their *in vivo* efficacy in relevant disease models.

## Results and discussion

### Inducible expression of MLKL and ASC intrabodies block necrotic cell death

We first validated the anti-necrotic capacity of two published intrabodies, a MLKL inhibitory monobody, Mb37_MLKL_ [11], and ASC targeting nanobody (VHH_mASC_) [32]. Mb37_MLKL_, and a control monobody Mb32_MLKL_ that binds to, but does not inhibit, MLKL [11], were FLAG-tagged at the N-terminus and fused with GFP at the C-terminus (**Fig. 1A**). The inhibitory VHH_mASC_[32] was also tested as a self-separating homodimer (VHH_mASC_-T2A-VHH_mASC_), with the two fused nanobodies modified with distinct HA and FLAG epitopes and split by a T2A sequence to enable their separation and detection, while VHH_NP-1_ [33], a nanobody targeting the nucleocapsid of influenza A virus, was used as a negative control (**Fig. 1A**). To assess intrabody expression, doxycycline (dox)-inducible stable HT29 cell lines containing MLKL intrabodies and immortalized bone marrow derived macrophages (iBMDMs) cells harbouring ASC intrabodies were generated. Following induction with dox, immunoblot analysis confirmed successful protein expression of all intrabody constructs at the expected molecular weight (**Fig. 1B, 1C**). Of note, the T2A motif containing ASC intrabody showed excellent separation into two monomeric nanobodies, with only a faint band corresponding to the full-length ASC intrabody fusion detected (**Fig. 1C**).

**Figure 1.**
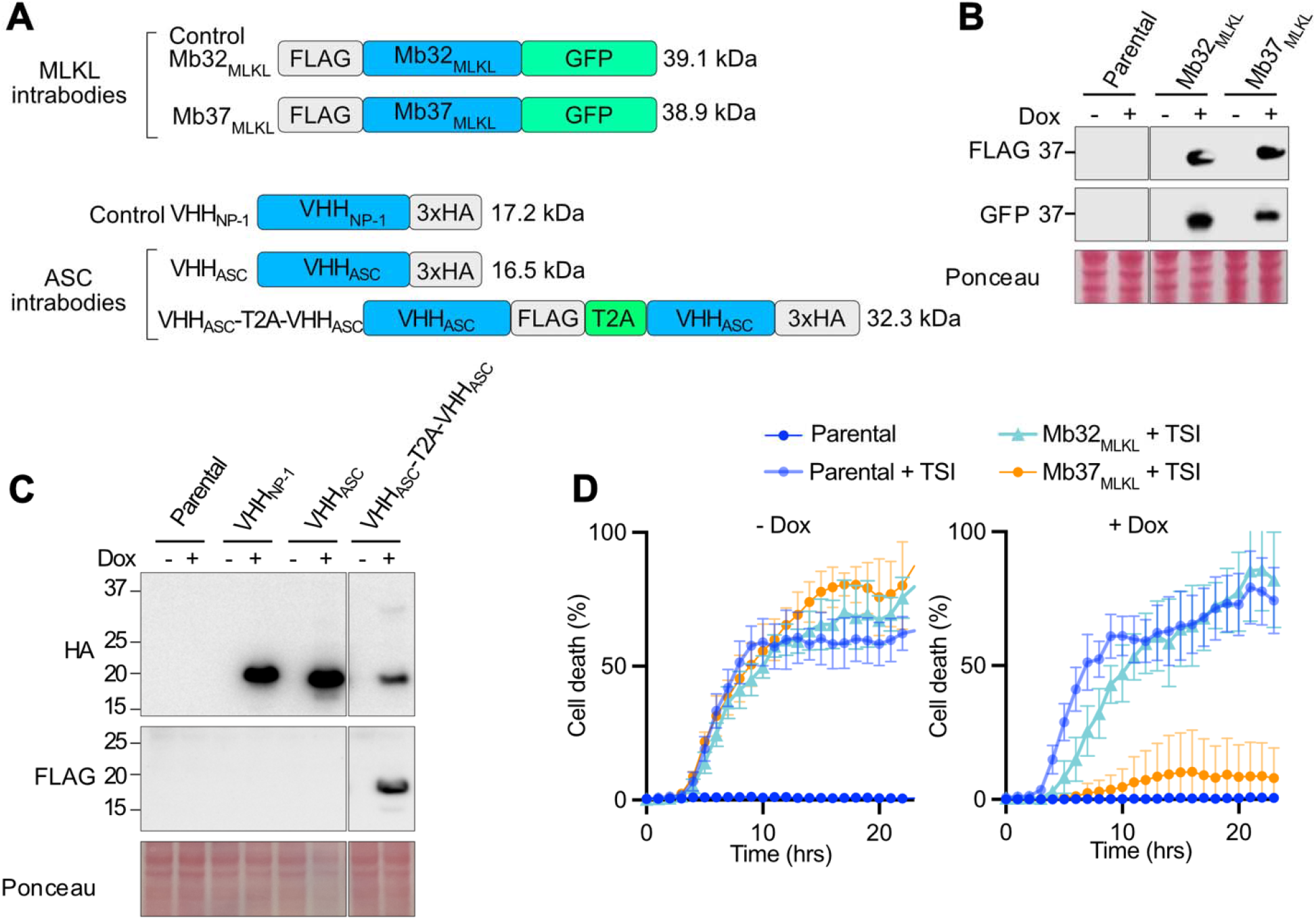
Doxycycline-inducible MLKL intrabody expression blocks necroptosis. **(A)** Schematic of doxycyline-inducible constructs. **(B)** Immunoblot analysis following overnight dox (100 ng/mL) treatment to induce expression of the indicated MLKL intrabodies in HT29 cells. **(C)** Immunoblot analysis following overnight dox (100 ng/mL) treatment to induce expression of ASC and NP-1 control intrabodies in iBMDMs. **(D)** Cell death kinetics of wildtype parental and stable HT29 cells following TSI (TNF (50□ng/mL), Smac mimetic Compound A (1□µM) and pan-caspase inhibitor IDN-6556 (10□µM)) stimulation with or without dox (0.1 µg/mL) pre-treatment overnight. Error bars represent standard deviation (SD) from three technical repeats. Data presented in B, C and D are representative of two independent experiments.

Next, the functional efficacy of MLKL targeting Mb37_MLKL_ was examined following overnight dox induction and subsequent treatment with the necroptotic stimulus, TSI: a combination of TNF, the Smac mimetic Compound A (Cp. A), and the pan-caspase inhibitor IDN-6556. Cell death analysis over time using an IncuCyte live cell imaging system demonstrated that, as expected [11], Mb37_MLKL_ expression abolished necroptotic cell death for more than 20 hours post TSI treatment, while expression of the control intrabody, Mb32_MLKL_, did not (**Fig. 1D**).

Pyroptosis and its ability to be inhibited by the ASC targeting intrabody was similarly evaluated by pre-treating iBMDM stable cell lines with dox overnight followed by nigericin stimulation to activate the NLRP3 inflammasome. Expression of monovalent VHH_mASC_ or VHH_mASC_-T2A-VHH_mASC_ intrabody protected cells from NLRP3-driven pyroptosis, while control VHH_NP-1_ intrabody-expressing cells, or wildtype parental iBMDMs, were not protected from nigericin killing (**Fig. 2A and Fig. S1**). Notably, the VHH_mASC_-T2A-VHH_mASC_ intrabody exhibited stronger protection from pyroptosis than its monovalent counterpart and was comparable to the protection resulting from MCC950 treatment (**Fig. 2A**), a selective and potent NLRP3 inhibitor [34]. Moreover, while MCC950 treatment conferred additional protection from nigericin-induced pyroptosis when used together with expression of the monovalent VHH_mASC_ intrabody (**Fig. 2A**), it did not increase protection substantially when used in conjunction with the VHH_mASC_-T2A-VHH_mASC_ intrabody (**Fig. 2A**). This indicates that the VHH_mASC_-T2A-VHH_mASC_ intrabody likely confers near-maximal protection from NLRP3 signalling, and that the nigericin-induced death observed after ∼ 5 hours of treatment in the presence of VHH_mASC_-T2A-VHH_mASC_ intrabody or MCC950 reflects NLRP3-independent killing. Therefore, while monovalent Mb37 intrabody suffices to shut down necroposis signalling, a VHH_mASC_-T2A-VHH_mASC_ intrabody is required to efficiently inhibit pyroptotic cell death.

**Figure 2.**
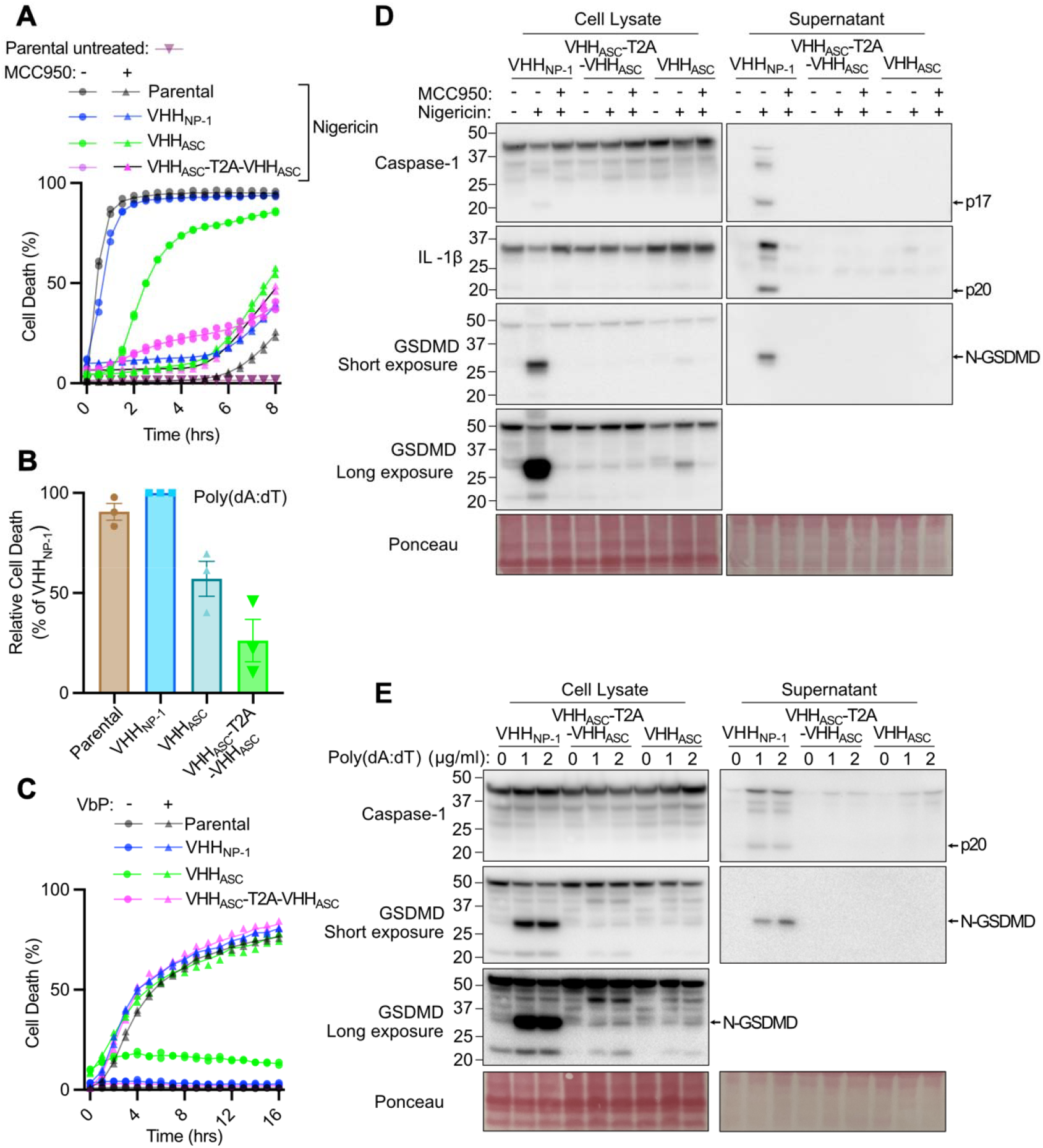
Doxycycline-inducible ASC intrabody expression prevents NLRP3- and AIM2-induced pyroptosis. (**A, B and C**) Wildtype parental and the indicated stable iBMDMs were pre-treated overnight with doxycycline (0.1 µg/mL) and cell death kinetics were monitored using IncuCyte imaging: (A) following stimulation with nigericin (5 µM) with or without MCC950 (10 µM) co-treatment; (B) after poly(dA:dT) (1 µg/ml) transfection; and (C) after VbP (2.5 µM) treatment. (**D and E**) Immunoblot analysis of pyroptotic signaling in total cell lysates and supernatants after (D) nigericin (5µM) ± MCC950 (10 µM) and (E) poly(dA:dT) treatment (1 and 2 µg/mL). In (B) data are presented as mean ± SEM of pooled data from three independent experiments. The IncuCyte data (A and C) symbols represent technical duplicates, and these experiements and the immunoblots (D and E) are representative of two independent experiments.

To further confirm the efficiency and specificity of monovalent and VHH_mASC_-T2A-VHH_mASC_ intrabodies, iBMDMs were treated with poly(dA:dT) to activate the AIM2 inflammasome that also requires ASC for pyroptotic killing, or were stimulated with Val-boroPro (VbP), which triggers NLRP1-mediated pyroptosis that occurs independent of ASC in mouse cells [35]. Consistent with ASC intrabody blockade of NLRP3 killing, VHH_mASC_-T2A-VHH_mASC_ intrabody expression conferred optimal protection from AIM2-mediated pyroptosis when compared to the monovalent ASC intrabody (**Fig. 2B**) but, as expected, neither ASC intrabody protected against NLRP1-mediated pyroptosis (**Fig. 2C**).

Consistent with the analysis of NLRP3- and AIM2-driven cell death, immunoblots showed that NLRP3- and AIM2-induced processing (*i*.*e*., activation) of caspase-1, IL-1β and Gasdermin D (GSDMD) was effectively shut down by ASC intrabody expression. The VHH_mASC_-T2A-VHH_mASC_ intrabody also better protected from GSDMD cleavage into its pore-forming N-terminal fragment (N-GSDMD) compared to VHH_mASC_ and acted similar to MCC950 treatment in the context of nigericin-induced NLRP3 triggering (**Fig. 2D and 2E**). Together, these results demonstrate that the inducible expression of MLKL and ASC targeting intrabodies can effectively shut down necrotic cell death signalling.

### Efficient MLKL intrabody delivery, expression and inhibition of necroptosis using LNP-mRNA technology

Following our validation and optimization of anti-necrotic intrabodies using a stable cell line dox inducible system (**Fig. 1 and Fig. 2**) Mb32_MLKL_ control and Mb37_MLKL_ were cloned into an *in vitro transcription* (IVT) vector incorporating a optimized synthetic 5′ UTR, a 2x human β-globin 3′ UTR, and a poly(A) tail (≥128 A bases) [36] and mRNA synthesized. LNPs containing mRNA were formulated using SM-102 (ionizable lipid), 1,2-distearoyl-sn-glycero-3-phosphocholine (DSPC), cholesterol, and 1,2-Dimyristoyl-sn-glycero-3-methoxypolyethylene glycol (DMG-PEG) at a 50:10:38.5:1.5 molar ratio and encapsulated using a NanoAssemblr Ignite microfluidic system at an N/P ratio of 6. The physiochemical characterisation of LNPs and mRNA loading was performed by dynamic light scattering and Qubit analysis. This showed that LNPs were ∼100 nm in diameter with unimodal distribution and poly dispersity index (PDI) <0.2, and no less than 85% mRNA encapsulation efficiency (**Fig. S2 and Table S1**). Next, LNPs containing FLAG-Mb32_MLKL_-GFP and FLAG-Mb37_MLKL_-GFP mRNA were examined for their transfection efficiency in three necroptosis susceptible cell lines (HT29, Colo205, SW620) using GFP as a readout and measured by widefield fluorescence microscopy and IncuCyte live cell imaging. Robust GFP expre ssion was observed across all cell lines, reaching ∼100% within 12 hours of LNP-mRNA incubation (**Fig. 3A and Fig. S3A, S3B**). Consistent with this data, immunoblotting showed an LNP-mRNA dose-dependent expression of FLAG-Mb32_MLKL_-GFP and FLAG-Mb37_MLKL_-GFP fusion proteins at their predicated molecular weights (**Fig. 3B**). The intracellular stability of LNP-delivered MLKL intrabody protein was assessed by treating cells with the protein synthesis inhibitor cycloheximide. While the short-lived protein Mcl-1 rapidly degraded within four hours of cycloheximide incubation, MLKL intrabody protein remained stable for at least eight hours across all cell lines (**Fig. 3C**).

**Figure 3.**
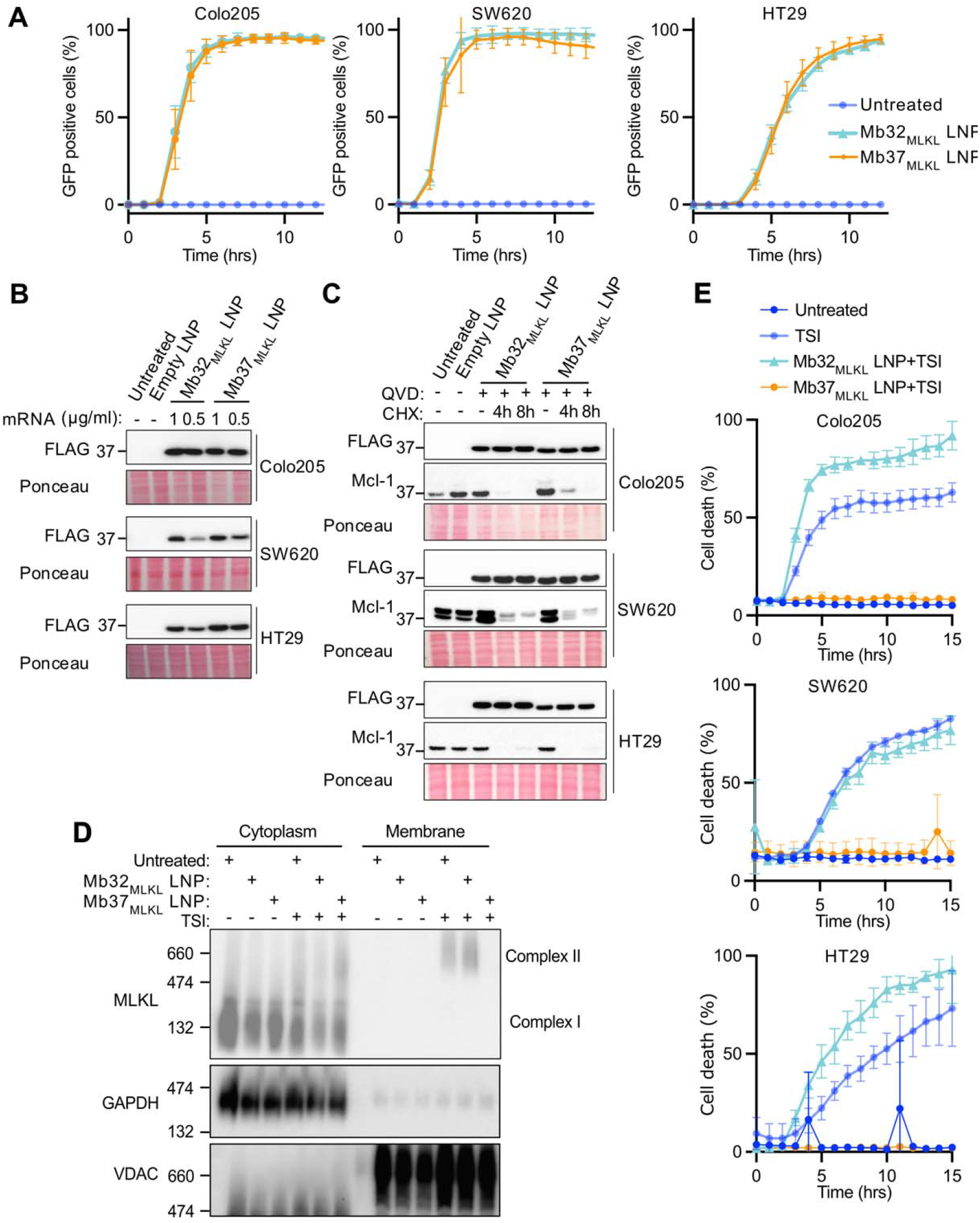
LNP-delivered MLKL intrabody mRNA prevents necroptosis by blocking MLKL membrane translocation. **(A)** Percentage of GFP-positive cells over time following LNP delivery of FLAG-MLKL-GFP intrabody mRNA (0.5 µg/mL) in Colo205, SW620, and HT-29 cells, measured using IncuCyte live cell imaging system. Error bars represent the SD of three technical replicates. See Figure S3 for representative images. **(B)** Immunoblot of total cell lysates 24 hours post-LNP transfection using the indicated mRNA concentrations. **(C)** Immunoblot of total lysates from cells treated with LNPs (1 µg/mL mRNA equivalent) for 24 hours, followed by cycloheximide (CHX, 40 µg/mL) and Q-VD-OPh (QVD, 20 µM) treatment as indicated. **(D)** Analysis of MLKL oligomerization and membrane translocation via Blue Native PAGE after LNP delivery of the indicated MLKL intrabody mRNA (0.5 µg/mL mRNA equivalent) overnight followed by TSI (TNF (50□ng/mL), Smac mimetic Compound A (1□µM), pan-caspase inhibitor IDN-6556 (10□µM)) treatment for 6 hours. GAPDH and VDAC were used as cytoplasmic and membrane markers, respectively. **(E)** Cell death kinetics measured by IncuCyte imaging following TSI (TNF (50□ng/mL), Smac mimetic Compound A (1□µM), pan-caspase inhibitor IDN-6556 (10□µM)) treatment in cells that were pretreated overnight with LNPs (0.5 µg/mL mRNA equivalent). Error bars represent SD from three technical replicates. All IncuCyte data (A, E) and immunoblots (B, C, D) are representative of two independent experiments.

Next, the function of LNP delivered MLKL intrabody was evaluated. Blue Native-PAGE analysis demonstrated that LNP delivery of Mb37_MLKL_, but not Mb32_MLKL_, blocked MLKL membrane translocation (**Fig. 3D**), consistent with our previous data showing that Mb37_MLKL_ inhibits MLKL function downstream of oligomerization by preventing MLKL membrane association [11]. In addition, like our dox inducible cell line data (**Fig. 1 and 2**), LNP delivered Mb37_MLKL_ completely shut down necroptotic cell death in HT29, Colo205 and SW620 cells, while control Mb32_MLKL_ did not (**Fig. 3E**). Importantly, LNP-delivered MLKL intrabodies did not block apoptosis induced by the BH3-mimetic ABT737 and S63845 treatment (**Fig. S4**).

### LNP delivery ASC intrabody mRNA blocks pyroptotic responses

Our stable doxycycline inducible ASC intrabodies showed that VHH_mASC_-T2A-VHH_mASC_ was superior at inhibiting pyroptosis when c ompared to a single VHH_mASC_ (**Fig. 2**). We therefore generated LNPs containing both self-separating VHH_mASC_-T2A-VHH_mASC_ mRNA and, as a comparison, non-cleavable bivalent VHH_mASC_-GS-VHH_mASC_ mRNA (**Fig. 4A, Fig. S2 and Table S1**). iBMDMs treated with LNP-mRNAs showed robust ASC intrabody protein expression, including excellent separation of VHH_mASC_-T2A-VHH_mASC_ into monomeric intrabodies (**Fig. 4B**). By comparison, however, VHH_NP-1_-T2A-VHH_NP-1_ relatively poor and hard to detect (**Fig. 4B**). Regardless, both bivalent and self-separating ASC intrabodies, but not control VHH_NP-1_-T2A-VHH_NP-1_, protected iBMDMs from nigericin-induced pyroptosis comparably, and to a similar extent as MCC950 (**Fig. 4C**). LNP delivered bivalent ASC intrabody mRNA also protected cells from AIM2-induced pyroptosis (**Fig. 4D**). Consistent with these data, and akin to dox inducible ASC intrabody expression (**Fig. 2D and 2E**), LNP delivered bivalent ASC intrabody mRNA sufficed to inhibit NLRP3- and AIM2-driven caspase-1, IL-1β and GSDMD activation, as reflected by the inhibition of processing into their p20, p17 and pore-forming N-GDSMD fragments, respectively (**Fig. 4E and F**).

**Figure 4.**
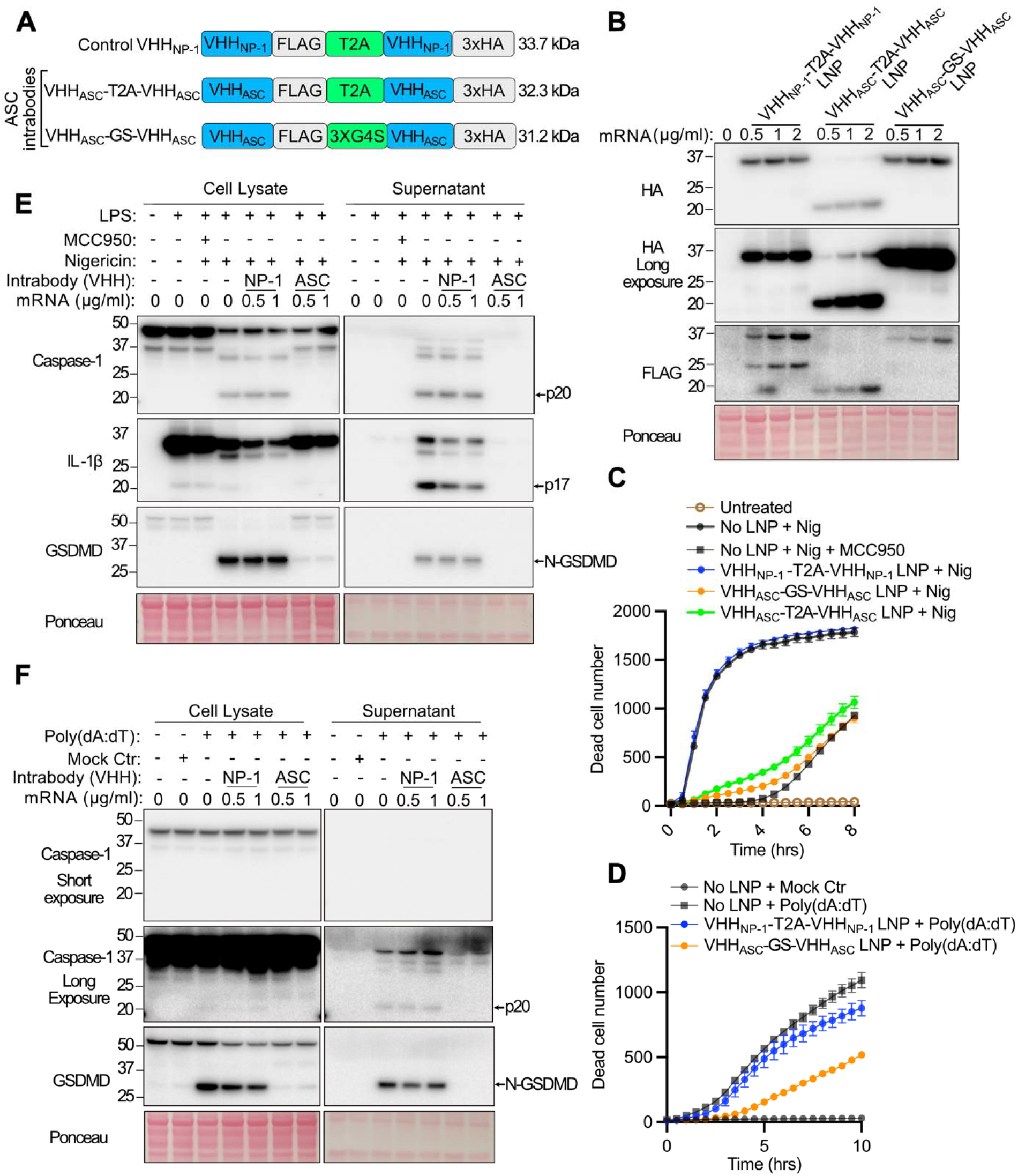
LNP-delivered ASC intrabody mRNA limits pyroptosis. **(A)** A schematic of mRNA constructs. **(B)** Immunoblot analysis of whole-cell lysates harvested after LNP-mediated treatment overnight at the specified mRNA concentrations followed by nigericin (5 µM) stimulation for 1 hour. (**C and D**) iBMDMs were treated overnight with LNPs (0.5 µg/mL mRNA equivalent) and then stimulated with (C) nigericin (5 µM) with our without MCC950 (10 µM), or transfected with (D) poly(dA:dT) (1 µg/mL). Cell death was measured by IncuCyte imaging. Error bars represent the SD of three technical replicates. (**E and F**) Immunoblot analysis of total lysates from iBMDMs pre-treated with increasing concentrations of LNP-mRNA overnight followed by (E) LPS (50 ng/mL) priming for 3 hrs and then nigericin (5 µM) treatment for 1 hr with or without MCC950 (10 µM), or (F) poly(dA:dT) (1 µg/mL) transfection for 6 hrs. All data (B to F) are representative of two independent experiments.

## Conclusions

Overall, this study presents a clinically relevant, small molecule alternative, approach for inhibiting intracellular proteins. By deploying intrabody mRNA encapsulated in LNPs as agents to block key proteins required for cellular necrosis, MLKL and ASC, we demonstrate substantial inhibition of inflammatory cell death *in vitro*, and show that, in the case of ASC targeting, bivalent ASC intrabody, or self-separating ASC intrabody homodimers, exhibit superior protection against pyroptosis compared to monovalent ASC nanobody. Moreover, the functional parity between LNP-delivered intrabody mRNA and those expressed via a doxycycline inducible system underscores the robustness, and clinical promise, of this delivery modality. These findings therefore lay the groundwork for advancing LNP-encapsulated intrabody mRNAs into relevant pre-clinical necrotic animal disease models, potentially extending antibody therapeutics to intracellular targets. This novel approach may have benefits compared to conventional therapies such as anti-cytokine biologics and small molecule inhibitors, as these are often constrained by systemic side effects, high costs, and variable efficacy.

## Materials and Methods

### Cell culture

HT29 and HEK293T cells, as well as NLRP3-FLAG/ASC-mCherry expressing iBMDMs [37] were cultured in DMEM (Gibco, Cat# 11885-084) supplemented with 10% (v/v) fetal bovine serum (FBS; Sigma, Cat# F9423-500ML or Gibco, Cat# 10099-141) and antibiotics (100 U/mL penicillin, and 100 µg/mL streptomycin), at 37□°C in a humidified incubator with 10% CO□. Colo205 and SW620 cells were maintained in RPMI-1640 (Gibco, Cat# 31800089) supplemented with the 10% (v/v) FBS under identical conditions. While the cell lines were not formally authenticated via genetic sequencing, their morphologies and responses to stimuli were consistent with their reported identities.

### Plasmid construction and lentiviral generation of stable cell lines

Constructs were cloned into a dox-inducible pF TRE3G PGK puro plasmid [38] (**Table S2 and Table S3**). Constructs were either purchased from Genscript or assembled in-house using the NEBuilder HiFi DNA Assembly Cloning Kit (NEB, Cat# E5520S) and gBlocks Gene Fragments (IDT). Final plasmids were sequence-verified by Sanger sequencing. Full length plasmid sequences can be provided upon request.

Lentiviral particles were produced by co-transfecting HEK293T cells with packaging and envelope plasmids (pVSVg, pMDL, and pRSV-REV) using Lipofectamine 3000 (Thermo Fisher, Cat# L3000008). Viral supernatants were harvested at 48- and 72-hours post-transfection, filtered through a 0.45□µm syringe filter (Sartorius Cat# 6533K), and used to transduce target cells in the presence of 8□µg/mL polybrene (Sigma, Cat# 107689). Successfully transduced cells were selected with 1µg/ml of puromycin (InvivoGen, Cat# ant-pr-5).

### IVT plasmid and mRNA generation

Gene blocks were amplified via PCR using primers containing NcoI-HF and XhoI (NEB, Cat# R3193 and R0146) restriction sites (**Table S2**). PCR products were gel-purified and ligated into IVT plasmid pre-digested with NcoI and NotI. Ligation reactions were performed using T4 DNA ligase (Promega, Cat# M1801), and the constructs were transformed into DH10α competent *E*. *coli* by heat shock. Colonies were selected on ampicillin (100µg/ml)-containing agar plates, and clones positive for insert and polyA were validated by Sanger sequencing and restriction digest analysis with SapI and XbaI (NEB, Cat# R0569 and Cat# R0145), respectively. Midipreps (ZymoPURE II Plasmid Midiprep Kit, Cat# D4200) were performed for the correct clones to isolate plasmid DNA for in vitro transcription (IVT).

For IVT, plasmids (**Table S3**) were linearized using SapI (NEB, Cat# R0569) and purified using the QIAquick PCR Purification Kit (Qiagen Cat# 28104). Uncapped RNA was synthesized using the HiScribe T7 High Yield RNA Synthesis Kit (NEB, Cat# E2040), while capped RNA was synthesized using the HiScribe T7 mRNA Kit with CleanCap® Reagent AG (NEB, Cat# E2080). A one-pot Cap-1 structure synthesis was performed using Faustovirus capping enzyme and 2′-O-methyltransferase (Company, Cat# #M2081, #M0366). After each step, RNA was purified using the Monarch RNA Cleanup Kit (NEB, Cat# T2050). RNA quality was assessed using a Nanodrop, Qubit™ RNA broad range kit(Thermo Fisher, Cat# Q10210), and Tapestation (Agilent Cat# 5067-5576). Capped mRNA was stored at –80□°C until use.

### LNP synthesis

mRNA-loaded LNPs were synthesized using a one-step microfluidic mixing protocol on the NanoAssemblr Ignite system. Lipids—SM102 (ionizable lipid), DSPC (zwitterionic lipid), cholesterol, and DMG-PEG—were prepared as 10□mM solutions in ethanol and mixed in a molar ratio of 50:10:38.5:1.5. The mRNA was diluted in 50□mM sodium acetate buffer (pH 4.0) to achieve an aqueous-to-ethanol flow rate ratio of 3:1. The N/P ratio was set at 6, and the mixing was performed at a total flow rate of 12□mL/min. Synthesized LNPs were diluted 30-fold in DPBS (pH 7.4) and concentrated using 100□kDa centrifugal filters (Millipore, Cat# UFC9100) at 4,000□rpm for 20 minutes. Particle size and polydispersity index (PDI) were determined by dynamic light scattering (DLS) following dilution (5□µL in 45□µL PBS). RNA encapsulation efficiency was measured using the Qubit RNA Broad-Range Assay after treating LNPs with either TE buffer or TE containing 2% Triton X-100. The encapsulated RNA content was calculated by subtracting the untreated (TE) signal from the lysed (Triton-treated) signal, expressed as a percentage of the input RNA.

### Immunoblotting

Cells were seeded into 6- or 12-well tissue culture treated plates (Corning®, Cat #3506 and #3512) and incubated for 24 hours before dox induction (100□ng/mL) or treatment with LNPs containing 1µg/ml or 0.5µg/ml of mRNA equivalent, alone or in combination with other drugs, as specified in relevant figure legends. Cells were lysed in 2× SDS Laemmli buffer, heated at 95□°C for 10–15 minutes, and proteins separated using 4%–12% gradient Bis-Tris SDS-PAGE gels (Invitrogen, Cat# NP0321). Proteins were transferred onto nitrocellulose membranes (Amersham, Cat# GE10600073). Ponceau S staining was routinely performed to verify equal protein loading. Membranes were blocked with 5% (w/v) skim milk in TBS containing 0.1% Tween 20 (TBS-T) and probed overnight at 4□°C with primary antibodies in blocking buffer with 0.04% sodium azide. Horseradish peroxidase-conjugated secondary antibodies were incubated for 1 hour at room temperature. Blots were developed using ECL (Millipore, Bio-Rad) and imaged using the ChemiDoc Touch Imaging System (Bio-Rad). Antibody details are provided in the **Table S4**.

### IncuCyte cell death assays

Colo205, SW620, and HT29 cells were plated at 3,000 cells/well in 96-well plates and allowed to adhere for 24 hours. Cells were pre-treated with doxycycline (100□ng/mL) overnight prior to the addition of reagents, including TNF (50□ng/mL), Smac mimetic Compound A (1□µM), pan-caspase inhibitor IDN-6556 (10□µM), ABT737 (1□µM), and S63845 (10□µM), alone or in combination. For iBMDMs, cells were plated at 10,000 cells/well in 96-well plates and allowed to adhere for 24 hours. Cells were pre-treated with doxycycline (100□ng/mL) overnight prior to the addition of reagents including nigericin (5□µM, Sigma, Cat# N7143), or Val-boroPro (VbP, 2.5 µM, InvivoGen, Cat# tlrl-vbp), or transfection of poly(dA:dT) (InvivoGen, Cat# tlrl-patn) using Lipofectamine 3000. Cell death was assessed using propidium iodide (PI, 0.2□µg/mL), SYTOX Green (0.5□µM, Thermo Fisher and cat# S7020), or TO-PRO-3 Stain (Thermo Fisher and cat#T3605, NIR dead cell dye, 1:5000 dilution) combined with nuclear dyes SPY505, SPY595, or SPY700 (Spirochrome, Cat# SPY505-DNA, SPY595-DNA, or SPY700-DNA). Images were captured using the IncuCyte SX5 or S3 imaging system (software versions v2022B or v2021B). The percentage of cell death was calculated by dividing the number of TO-PRO-3, PI or SYTOX Green-positive cells (dead) by the total number of SPY-positive cells (live and dead).

## Acknowledgements

We thank Eicke Latz and Ashley Mansell for providing NLRP3-FLAG/ASC-mCherry expressing iBMDMs. This work was supported by National Health and Medical Research Council of Australia (NHMRC) Investigator Grants (2008692 to JEV; 1172929 and 2034104 to JMM), a mRNA Victoria Research Acceleration Fund Grant (to JEV, DG and HM) and operational infrastructure grants through the NHMRC Independent Research Institute Infrastructure Support Scheme (9000719) and the Victorian State Government Operational Infrastructure Support.

## Declarations of interests

JEV, CB and RS own shares in Mermaid Bio GmbH. JMM has received research funding from, and contributes to a project developing necroptosis pathway inhibitors in collaboration with, Anaxis Pharma Pty. Ltd.

