## Supplemental data for "Lipid nanoparticle delivered intrabodies for inhibiting necroptosis and pyroptosis"

**Supplemental Figures and Tables**

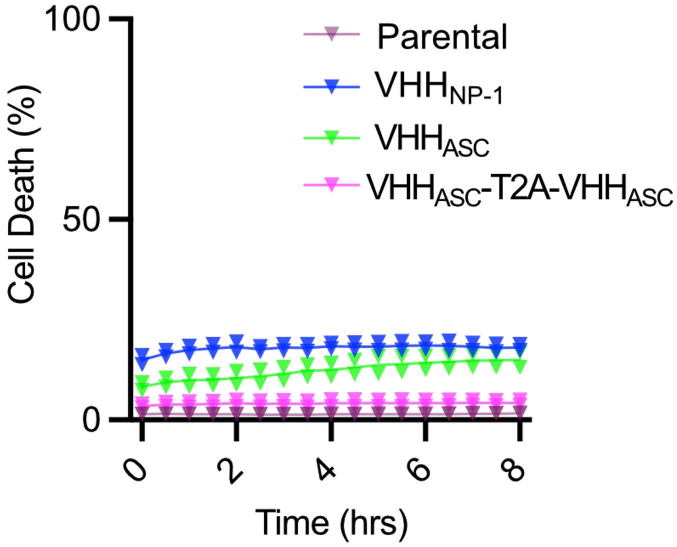


**Figure S1. Intrabody expression alone does not cause cell death.** Wildtype parental and the indicated stable iBMDM lines were treated overnight with 0.1 µg/mL doxycycline and cell death analysed. Represents control data (*i.e.,* no nigericin treatment) for Figure 2A.

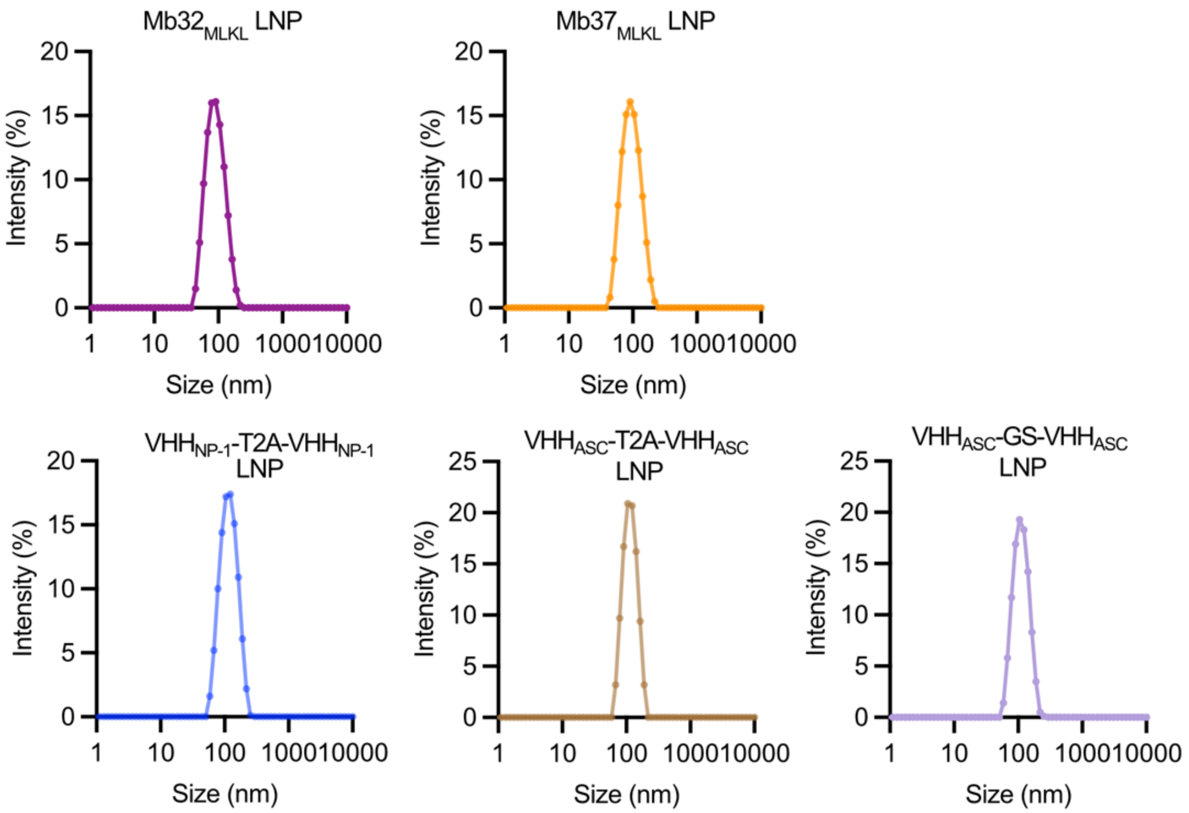


**Figure S2. Unimodal LNP-mRNA size.** Size distribution of LNP-mRNAs as measured by dynamic light scattering.

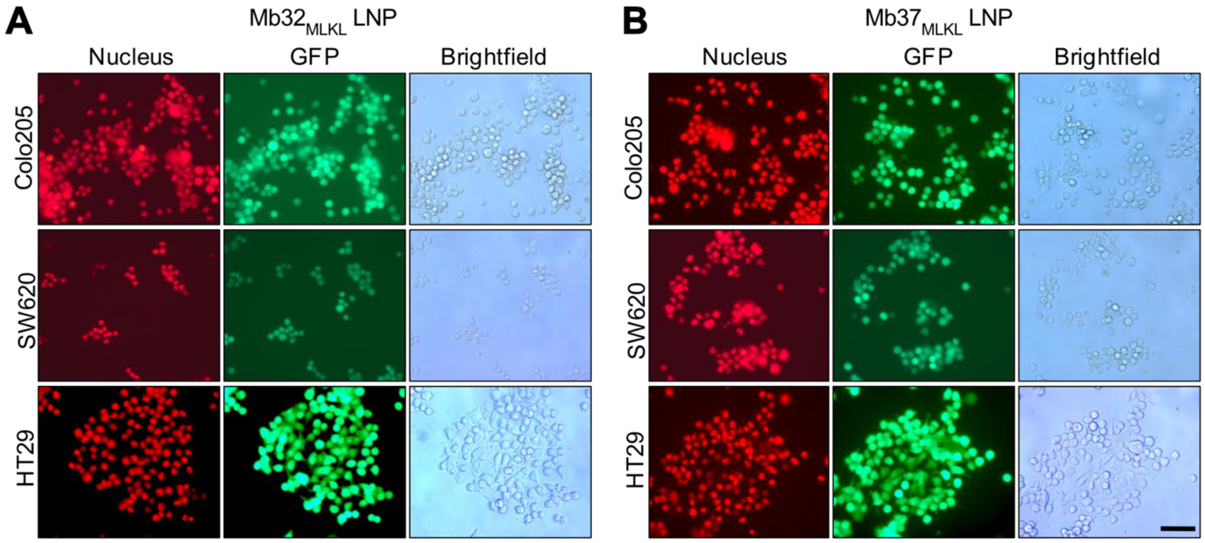


**Figure S3. LNP-delivered MLKL intrabodies are expressed in the majority of cells.** Widefield fluorescence microscopy recorded 24 hours after LNP-mRNA treatment 0.5 µg/mL mRNA equivalent in the indicated cancer cell lines. Nuclei were stained with SPY DNA 620. Scale bar, 200 µm.

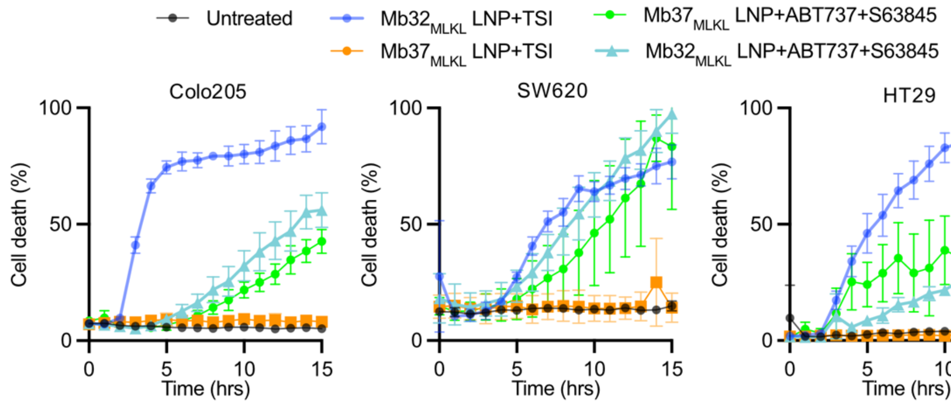


**Figure S4. MLKL intrabodies do not block intrinsic apoptosis.** Cell death kinetics measured by IncuCyte live-cell imaging following treatments to induce either necroptosis or apoptosis. Necroptosis was triggered using a TSI stimuli consisting of TNF (50 ng/mL), Smac mimetic Compound A (1 µM), and the pan-caspase inhibitor IDN-6556 (10 µM). Intrensic apoptosis was induced using ABT-737 (1 µM) and S63812 (5 µM). Prior to cell death stimuli, cells were pretreated overnight with $Mb32_{MLKL}$ LNP or $MB37_{MLKL}$ LNPs at 0.5 µg/mL mRNA equivalent. Error bars represent the standard deviation of three technical replicates. The data are representative of two independent experiments.

**Table S1. Physiochemical characterisation of LNP-mRNAs.**

| LNP | Z-average diameter (nm) | PDI | mRNA loading efficiency (%) |
|---|---|---|---|
| $Mb32_{MLKL}$ LNP | 88.7 | 0.13 | 87.4 |
| $Mb37_{MLKL}$ LNP | 85.6 | 0.11 | 89.2 |
| $VHH_{NP-1}$-T2A-$VHH_{NP-1}$ LNP | 106.0 | 0.054 | 85.6 |
| $VHH_{mASC}$-GS-$VHH_{mASC}$ LNP | 110.3 | 0.045 | 86.4 |
| $VHH_{mASC}$-T2A-$VHH_{mASC}$ LNP | 110.7 | 0.066 | 89.4 |

**Table S2. Sequence of primers and gene blocks.**

| Primer/gene block | Sequence |
|---|---|
| **Primer** | |
| Forward primer for FLAG-Mb32$_{MLKL}$-GFP and FLAG-Mb37$_{MLKL}$ -GFP | 5'-<br>TGACCCATGGTCTAGACCACCATGGATTACAAAGACGATG<br>ATGAT-3' |
| Reverse primer for FLAG-Mb32$_{MLKL}$ -GFP and FLAG-Mb37$_{MLKL}$ -GFP | 5'-TTATCTCGAGTTACTTGTACAGCTCGTCCATGCC-3' |
| **Gene block** | |
| FLAG-Mb32 $_{MLKL}$-GFP | 5'-CACCATGGATTACAAAGACGATGATGATAAGGGATCCGT<br>TTCTTCTGTTCCGACCAAACTGGAAGTTGTTGCTGCGACCC<br>CGACTAGCCTGCTGATCAGCTGGGATGCTCCTGCAGTTACC<br>GTCGATCTTTACATTATCACGTACGGTGAAACCGGTGGTAAC<br>TCCCCGGTTCAAACGTTTGAGGTACCAGGTTCCAAGTCTAC<br>TGCTACCATCAGCGGCCTGAGCCCGGGTGTCGACTATACCA<br>TCACTGTATACGCATACTCTTTCATGTACCATGACTACTACTA<br>CCCGGAATGGAGCCCAATCTCGATTAACTACCGTACCGCTA<br>GCAGTTCCTCTAGTGCGGCCGCTGTGAGCAAGGGCGAGGA<br>GCTGTTCACCGGGGTGGTGCCCATCCTGGTCGAGCTGGACG<br>GCGACGTAAACGGCCACAAGTTCAGCGTGTCCGGCGAGGG<br>CGAGGGCGATGCCACCTACGGCAAGCTGACCCTGAAGTTC<br>ATCTGCACCACCGGCAAGCTGCCCGTGCCCTGGCCCACCCT<br>CGTGACCACCCTGACCTACGGCGTGCAGTGCTTCAGCCGCT<br>ACCCCGACCACATGAAGCAGCACGACTTCTTCAAGTCCGC<br>CATGCCCGAAGGCTACGTCCAGGAGCGCACCATCTTCTTCA<br>AGGACGACGGCAACTACAAGACCCGCGCCGAGGTGAAGTT<br>CGAGGGCGACACCCTGGTGAACCGCATCGAGCTGAAGGGC<br>ATCGACTTCAAGGAGGACGGCAACATCCTGGGGCACAAGC<br>TGGAGTACAACTACAACAGCCACAACGTCTATATCATGGCC<br>GACAAGCAGAAGAACGGCATCAAGGTGAACTTCAAGATCC<br>GCCACAACATCGAGGACGGCAGCGTGCAGCTCGCCGACCA<br>CTACCAGCAGAACACCCCCATCGGCGACGGCCCCGTGCTG<br>CTGCCCGACAACCACTACCTGAGCACCCAGTCCGCCCTGA<br>GCAAAGACCCCAACGAGAAGCGCGATCACATGGTCCTGCT<br>GGAGTTCGTGACCGCCGCCGGGATCACTCTCGGCATGGAC<br>GAGCTGTACAAGTAACTC-3' |
| FLAG-Mb37$_{MLKL}$-GFP | 5'-CCATGGATTACAAAGACGATGATGATAAGGGATCCGTTTC<br>TTCTGTTCCGACCAAACTGGAAGTTGTTGCTGCGACCCCGA |

| | |
|---|---|
| | CTAGCCTGCTGATCAGCTGGGATGCTCCTGCAGTTACCGTC<br>GATTATTACTATATCACGTACGGTGAAACCGGTGCTTCTTTC<br>TACTTCTACCAGACATTCAAGGTACCTGGTTCCAAGTCTACT<br>GCTACCATCAGCGGCCTGAGCCCGGGTGTCGACTATACCAT<br>CACTGTATACGCATACGACTGGGGTGGTTACTTCATCGGTAA<br>CCCAATCTCGATTAACTACCGTACCGCTAGCAGTTCCTCTAG<br>TGCGGCCGCTGTGAGCAAGGGCGAGGAGCTGTTCACCGGG<br>GTGGTGCCCATCCTGGTCGAGCTGGACGGCGACGTAAACG<br>GCCACAAGTTCAGCGTGTCCGGCGAGGGCGAGGGCGATGC<br>CACCTACGGCAAGCTGACCCTGAAGTTCATCTGCACCACCG<br>GCAAGCTGCCCGTGCCCTGGCCCACCCTCGTGACCACCCTG<br>ACCTACGGCGTGCAGTGCTTCAGCCGCTACCCCGACCACAT<br>GAAGCAGCACGACTTCTTCAAGTCCGCCATGCCCGAAGGC<br>TACGTCCAGGAGCGCACCATCTTCTTCAAGGACGACGGCA<br>ACTACAAGACCCGCGCCGAGGTGAAGTTCGAGGGCGACAC<br>CCTGGTGAACCGCATCGAGCTGAAGGGCATCGACTTCAAG<br>GAGGACGGCAACATCCTGGGGCACAAGCTGGAGTACAACT<br>ACAACAGCCACAACGTCTATATCATGGCCGACAAGCAGAA<br>GAACGGCATCAAGGTGAACTTCAAGATCCGCCACAACATC<br>GAGGACGGCAGCGTGCAGCTCGCCGACCACTACCAGCAGA<br>ACACCCCCATCGGCGACGGCCCCGTGCTGCTGCCCGACAA<br>CCACTACCTGAGCACCCAGTCCGCCCTGAGCAAAGACCCC<br>AACGAGAAGCGCGATCACATGGTCCTGCTGGAGTTCGTGA<br>CCGCCGCCGGGATCACTCTCGGCATGGACGAGCTGTACAAG<br>TAACTC-3’ |
| VHH$_{mASC}$-T2A-VHH$_{mASC}$ | 5’-ATAACGCGTCCATTCGACACGCCACCATGGCTCAGGTGC<br>AGCTGGTGGAGACTGGGGGCGGAATGGTCCATCCCGGTGG<br>CAGCCTGCGCCTGAGCTGTGCAGCCAGTGGCTTTACTTTCA<br>GCGAGTACGGGATGACTTGGGTGAGACAGGCTCCCGGGAA<br>GGGCCCCGAATGGGTGTCCAGGATTAACAGCAGTGGCGGA<br>TATACCGTATACAGGGCCTCTGTGAAAGGACGGTTTACCGT<br>GAGCAGAGACAACGCCAAGAACACCCTATACCTTCAGATG<br>AATTCACTGAAGCCAGAGGACACAGCCCTGTATTACTGTGC<br>CCGGACCACTAACTGGGAGACCAGACTATCTCAGGGCACC<br>CAGGTTACTGTCAGCAGCGGAGGTGACTACAAGGATGATG<br>ACGACAAGGGAGGGGGCGGCTCAGGAGGGGGGGTAGTG<br>GTGGTGGTGGCTCTCAGGTCCAGCTTGTGGAAACAGGAGG<br>AGGCATGGTGCACCCTGGTGGGTCACTGCGTCTGTCCTGCG<br>CCGCCTCCGGGTTCACCTTCTCTGAGTATGGCATGACCTGG<br>GTAAGACAAGCACCTGGCAAAGGCCCTGAGTGGGTTTCCA<br>GAATCAATTCCTCCGGCGGTTACACAGTCTACAGAGCTAGT<br>GTGAAGGGACGCTTCACGGTGAGCAGAGACAATGCCAAAA<br>ACACATTGTACCTGCAGATGAACTCATTAAAGCCAGAAGAC<br>ACAGCTTTGTACTACTGCGCCAGGACCACAAACTGGGAAA<br>CCAGACTGAGCCAAGGCACACAGGTGACAGTAAGCAGTGG<br>CGGTTACCCTTATGATGTGCCTGACTATGCTGGCTACCCATAT |

| | |
|---|---|
| | GACGTGCCAGACTACGCAGGTTCCTACCCCTACGATGTTCC<br>TGATTATGCCTGAGCGGCCGCTCGAGGCTCGCTTTCTTGCT<br>GTC-3’ |
| $VHH_{NP1}$-T2A-$VHH_{NP1}$ | 5’ATAACGCGTCCATTCGACACGCCACCATGGCTCAAGTACA<br>GCTCCAGGAAAGCGGTGGTGGACTCGTTCAAGCCGGGGGG<br>TCTCTGCGCCTTACTTGCGCACTTTCCGAGCGAACCAGCAC<br>TTCATACGCCCAGGGGTGGTTCAGACAGCCTCCCGGCAAG<br>GAGAGAGAATTCGTCGCAAGCCTCCGAACACATGATGGTA<br>ACACACACTATACCGATTCAGTGAAAGGTCGGTTTACAATAT<br>CTAGGGATAATGCCGAAAATACATTGTATCTCCAGATGAACT<br>CTCTTAAAACAGAAGATACAGCTGTGTACTACTGTGCCGCT<br>TCTCTCGGTTACTCAGGTGCTTACGCAAGCGGGTATGACTA<br>CTGGGGCCAAGGGACACAAGTTACAGTAAGCAGCGGTGGA<br>GATTACAAAGACGACGATGACAAGGGATCAGGCGAAGGCC<br>GGGGCAGCCTGCTCACCTGCGGAGATGTAGAAGAGAACCC<br>TGGGCCCCAGGTACAGCTGCAGGAATCAGGAGGGGGCCTG<br>GTGCAGGCTGGAGGGTCCCTTAGGCTGACATGCGCCCTCTC<br>TGAGAGGACCAGTACCTCATACGCTCAGGGCTGGTTTAGAC<br>AGCCACCCGGGAAGGAACGCGAGTTTGTCGCTTCCCTGCG<br>CACCCACGACGGAAATACTCATTATACAGATAGTGTTAAGG<br>GTCGGTTCACCATCAGTAGGGATAACGCTGAAAATACATTG<br>TACCTTCAAATGAATTCACTCAAGACCGAAGATACAGCAGT<br>CTACTATTGCGCAGCTAGTTTGGGTTACAGCGGAGCTTATGC<br>AAGTGGTTATGATTACTGGGGGCAGGGAACTCAGGTTACTG<br>TATCCAGTGGGGGTTATCCATATGATGTTCCAGACTATGCAG<br>GATATCCATACGATGTGCCCGACTACGCCGGCTCCTATCCAT<br>ACGACGTGCCAGATTATGCCTGAGCGGCCGCTCGAGGCTCG<br>CTTTCTTGCTGTC-3’ |

**Table S3. Plasmids used to make stable cell lines and IVT templates.**

| Plasmid | Description | reference |
|---|---|---|
| Doxycycline inducible plasmids | | |
| pF TRE3G_FLAG-MB32$_{MLKL}$-GFP<br>pF TRE3G_FLAG-MB37$_{MLKL}$-GFP | MLKL inhibitory monobody (Mb37) or control non-inhibitory monobody (Mb32) | Petrie, E. J. *et al*, 2020[1]. |
| pF_TRE3G_VHH$_{NP-1}$-3xHA | Control NP-1 nanobody | This work, Genscript |
| pF TRE3G_VHH$_{mASC}$-3xHA | ASC nanobody | This work, Genscript |
| pF TRE3G VHH FLAG-VHH$_{mASC}$-T2A_VHHm$_{ASC}$-3xHA | Self-separating ASC nanobody | This work, Genscript |
| IVT plasmids | | |
| pGEMs_eGFP_IVaxT_Poly_A_148 | Parental IVT plasmid | Fernández, S.L. *et*, 2021[2]. |
| pGEMs_VHH$_{NP-1}$-FLAG-T2A-VHH $_{NP-1}$-3xHA_IVT CleanCap | Control self-separating NP-1 nanobody | This work, made in house |
| pGEMs_VHH$_{mASC}$-FLAG-T2A-VHH$_{mASC}$-3xHA_IVT CleanCap | Self-separating ASC nanobody | This work, made in house |
| pGEMs_VHH$_{mASC}$-FLAG-3xG4S-VHH$_{mASC}$-3xHA_IVT CleanCap | Bivalent ASC nanobody | This work, made in house |
| pGEMs_ FLAG-MB32$_{MLKL}$-GFP_IVT | Control MLKL monobody | This work, made in house |
| pGEMs_ FLAG-MB37$_{MLKL}$-GFP_IVT | Inhibitory MLKL monobody | This work, made in house |


1 Petrie, E. J., Birkinshaw, R. W., Koide, A., Denbaum, E., Hildebrand, J. M., Garnish, S. E., Davies, K. A., Sandow, J. J., Samson, A. L., Gavin, X., Fitzgibbon, C., Young, S. N., Hennessy, P. J., Smith, P. P. C., Webb, A. I., Czabotar, P. E., Koide, S. and Murphy, J. M. (2020) Identification of MLKL membrane translocation as a checkpoint in necroptotic cell death using Monobodies. Proceedings of the National Academy of Sciences. **117**, 8468-8475

2 Linares-Fernandez, S., Moreno, J., Lambert, E., Mercier-Gouy, P., Vachez, L., Verrier, B. and Exposito, J. Y. (2021) Combining an optimized mRNA template with a double purification process allows strong expression of in vitro transcribed mRNA. Mol Ther Nucleic Acids. **26**, 945-956

**Table S4. Antibodies.**

| **Antibody** | **Company/catalogue number** | **Dilution** |
|---|---|---|
| Anti-FLAG M2-Peroxidase (HRP) antibody | Sigma-Aldrich/Cat# A8592 | 1:4000 |
| Rat anti-MLKL | Inhouse;<br>available from Merck Millipore / Cat# MABC604 | 1:1000 |
| Rabbit anti-MLKL (phospho S358) | Abcam/ Cat# ab187091 | 1:1000 |
| Rabbit anti-MCL-1 | Cell Signaling Technology/Cat# 5453 | 1:1000 |
| Rabbit anti-GFP | Invitrogen/Cat# A-6455 | 1:2000 |
| Mouse anti-VDAC1 | Merck Millipore/MABN504 | 1:500 |
| Rabbit anti-GAPDH | Cell Signaling Technology/Cat# 2118 | 1:1000 |
| Rabbit anti-ASC | Adipogen/Cat# AL177 | 1:1000 |
| Mouse anti-NLRP3 | Adipogen/ AG-20B-0014-C100 | 1:1000 |
| Mouse anti-IL1β | R&D/ Cat# AF-401-NA | 1:1000 |
| Mouse anti-Caspase 1 | Adipogen/ Cat# AG-20B-0042-C100 | 1:1000 |
| Rabbit anti-Caspase 1 | Abcam/ Cat# ab179515 | 1:1000 |
| Rabbit anti-GSDMD | Abcam/ Cat# ab209845 | 1:1000 |
| HRP-conjugated anti-rabbit secondary | Southern Biotech/ Cat# 4010-05 | 1:10000 |
| HRP-conjugated anti-mouse secondary | Southern Biotech/ Cat# 1010-05 | 1:10000 |
| HRP-conjugated anti-rat secondary | Southern Biotech/ Cat# 3010-05 | 1:10000 |
